# Sex-specific weighting of shoal size and movement speed but no evidence of asymmetric dominance effect in zebrafish shoal-size preference

**DOI:** 10.64898/2026.05.07.723409

**Authors:** Abhishek Singh, Neha Mary Mathew, Asmi Aggarwal, Tamanna Ail, Siya Kohli, Bittu Kaveri Rajaraman

## Abstract

Social decisions often require animals to integrate information across multiple attributes of potential partners. Using biological motion stimuli, point-displays generated from tracked live shoals, we tested how adult zebrafish (*Danio rerio*) weigh shoal size and movement speed during social preference, and whether these preferences are susceptible to contextual manipulation by an asymmetrically placed alternative. In Experiment 1, we established a multi-attribute indifference point by presenting males and females with dichotomous contrasts in which shoal size and movement speed were traded off. Both sexes showed no preference when a larger, slower shoal (4 fish at 0.75x speed) was pitted against a smaller, faster shoal (2 fish at 1.25x speed), but preferred the smaller, faster shoal when the speed difference was greater (4 fish at 0.5x versus 2 fish at 1.25x), indicating that zebrafish are sensitive to graded differences in movement speed relative to numerical cues. In Experiment 2, unidimensional tests confirmed that both sexes preferred larger shoals when speed was held constant but revealed sex-based differences in speed sensitivity: males preferred faster-moving shoals at both shoal sizes tested, whereas females showed no significant speed preference. Male shoal size preferences were stronger at higher movement speeds, suggesting that speed modulates the strength of size preference. In Experiment 3, we tested the asymmetric dominance effect in males, the only sex sensitive to both dimensions, using the indifferent contrast from Experiment 1 as the primary options and four decoy shoals asymmetrically placed along either the size or speed dimension, under counterbalanced presentation orders. No decoy shifted male preference significantly from chance under any condition. These results indicate that zebrafish weigh social cues in a sex-specific manner, and that asymmetric decoy options do not induce preference biases in males when shoals vary along the dimensions of movement speed and size.

## Introduction

The value of a choice option often depends on the context in which it is encountered rather than on its intrinsic properties alone (Huber et al., 1982; Tversky & Simonson, 1993). Classical normative models of decision-making, including expected utility and optimal foraging frameworks, assume that individuals assign stable, context-independent values to options and select those that maximize biological fitness (Stephens & Krebs, 1986; Kacelnik, 2006). A significant body of empirical work across humans and non-human animals indicates that preferences are frequently context-dependent, violating the assumption of value independence (Bateson, 2002; Latty & Trueblood, 2020; Spektor et al., 2021). One well-characterized example is the asymmetric dominance effect, in which two focal options trade off across attribute dimensions such that neither strictly dominates the other, and the introduction of a third option (the decoy) positioned asymmetrically along one of these dimensions can systematically bias preference between the primary alternatives (Huber et al., 1982; Latty & Beekman, 2011; Hemingway et al., 2024). By disproportionately influencing the relative weighting of one attribute dimension, such a decoy can shift preferences in ways that reflect context-dependent interactions between the psychophysical valuation of each attribute (Bateson et al., 2002, 2003; Latty & Beekman, 2011; Morgan et al., 2012; Rivalan et al., 2017; Latty & Trueblood, 2020).

While such violations are considered economically irrational, they may nonetheless reflect heuristic decision rules that are ecologically rational, allowing animals to efficiently navigate environments where the relative value of options varies across time and space (Gigerenzer et al., 2000; De Petrillo & Rosati, 2019; Kacelnik, 2006). Context-dependent preferences have been examined along single-attribute dimensions, where introducing a third option along the same dimension is expected to shift preferences between two primary alternatives (Morgan et al., 2012; Reding & Cummings, 2019; Singh et al., 2026). However, these studies differ fundamentally from tests of the asymmetric dominance effect in its canonical form, where a decoy is strategically positioned to shift preference between two primary options that trade off across two attribute dimensions simultaneously. This distinction is important because the asymmetric dominance effect has been argued to be qualitatively distinct from other forms of context-dependent bias, and its existence remains debated in the literature (Spektor et al., 2021; Armand et al., 2025).

Shoal size preference is well documented across fish species including zebrafish, with individuals consistently favouring larger groups that offer fitness benefits including reduced predation risk, improved foraging success, and increased reproductive opportunities (Svensson et al. 2000; Binoy & Thomas, 2004; Agrillo & Dadda, 2007; Buckingham et al. 2007; Seguin & Gerlai, 2017). Zebrafish social behaviour is strongly guided by visual information, encompassing both the physical appearance and motion patterns of conspecifics (Engeszer et al. 2008). Zebrafish have been shown to extract meaningful social information from movement cues alone, as experiments using robotic conspecifics and abstract motion stimuli where dot/s motion replicate the characteristic conspecific locomotor patterns (biological motion), demonstrate that dynamic visual information is sufficient to elicit affiliative behaviour even in the absence of recognisable morphological features (Larsch & Baier, 2018; Nunes et al. 2020).

Both point light displays (light points on a dark background) and point displays (dark points on a light background) have been used in single-choice and two-choice forced-preference paradigms to assess the discriminability of biological from non-biological motion and the capacity of such stimuli to elicit affiliative social behaviour (Schluessel et al. 2015, 2018; Nunes et al. 2020), with biological motion perception documented across multiple fish species—including zebrafish, *Danionella*, medaka, cichlids, and damselfish—suggesting it is a broadly conserved capacity among teleosts (Nunes et al. 2020; Larsch & Baier 2018; Zada et al. 2024; Nakayasu & Watanabe 2014; Schluessel et al. 2015, 2018). Larsch and Baier (2018) showed that zebrafish readily shoal with a single moving dot when it reproduces the bout kinematics of conspecific swimming, without requiring fish-like form, colour, or pigmentation pattern. Preferences for specific kinematic parameters tracked the animal’s own age-specific motion and were present even in isolation-reared individuals, suggesting an innate mechanism for biological motion recognition.

Yet the potential of biological motion stimuli extends beyond discriminability and affiliation, offering a tractable means to dissect the quantitative and kinematic attributes that drive shoaling decisions. Multi-dot displays that simulate shoal-like movement are particularly promising, as the shoal size (number of dots) and kinematics of dots can be independently controlled to represent ecologically relevant shoal properties. Here, we used multi-animal tracking of real shoals to generate biological motion stimuli in which shoal size and swimming speed could be varied both independently and presented in a trade-off manner. Using a standard two-option shoal preference task, we first established conditions under which fish showed no baseline preference between two shoals trading off across size and speed, while confirming sensitivity to each dimension in isolation. We then tested whether the introduction of a third option, asymmetrically placed along either the size or speed dimension, could bias preference toward one of the two otherwise indifferent shoals, providing a test of the asymmetric dominance effect in a naturalistic multiattribute decision context.

## Methods

### Subjects

Adult captive-bred zebrafish (*Danio rerio*; age < 1 year) were obtained from a local pet store in Daryaganj, New Delhi, India. Fish were housed in a ZebTec Active Blue Stand-Alone system (Tecniplast, PA, USA) at Ashoka University, Sonipat, Haryana, India. They were maintained under a 12:12 h light–dark cycle (lights on from 10:00 a.m. to 10:00 p.m.), with water parameters kept at 28–30 °C, pH 7.5–8.5, and conductivity between 650–700 µS. Fish were fed ad libitum twice daily with powdered Tetra-Tetramin flakes. A total of 419 individuals were used across all experiments (310 males; 109 females).

### Shoal choice apparatus

The shoaling apparatus consisted of a transparent cylindrical acrylic focal tank (40 cm diameter, 0.5 cm thickness) placed at the center of a larger square glass tank (50 × 50 × 29 cm, L × B × H; 0.5 cm thickness; Figure 1 B). Four display screens (Acer EK220Q 54.6 cm, 100Hz refresh rate) of dimensions 50 × 28 cm were positioned facing the four walls of the tank (Figure 1 B), at 4.5 cm from the focal cylindrical tank. All joints were sealed using silicone sealant.

**Figure 1.**
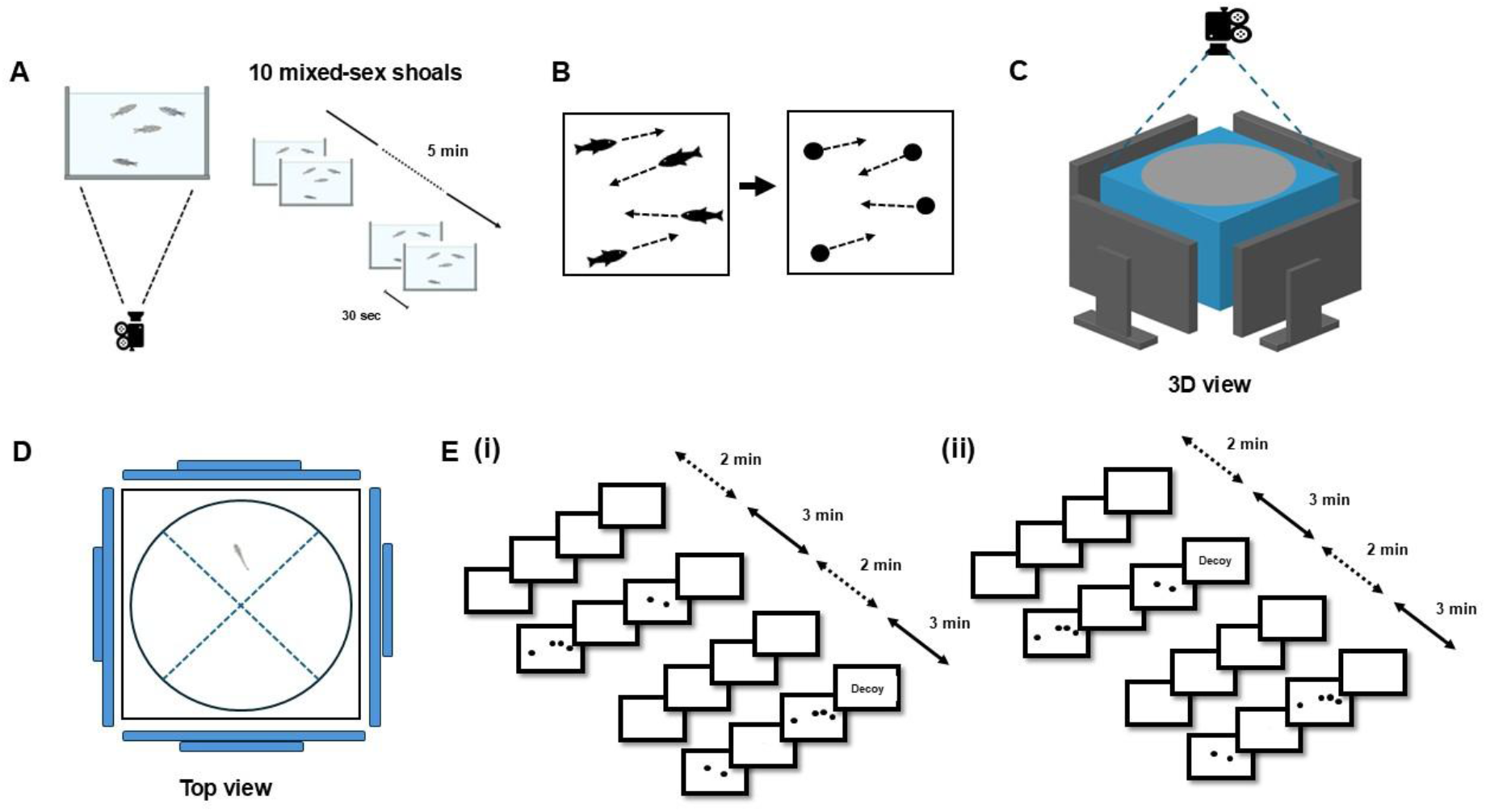
Virtual shoal generation, behavioural setup, and experimental design. **A.** Preparation and recording of stimulus shoals. Ten mixed-sex shoals were filmed, and 30 s trajectory segments were sampled from these recordings to generate 5 min stimulus movies. **B**. Trajectory extraction and animation. Individual fish were tracked to obtain movement trajectories, each represented as a moving point. Different-sized shoals were constructed from the same four-fish trajectory pool by randomly combining 30 s trajectory segments. **C**. Three-dimensional schematic of the behavioural setup showing the square tank with a central circular arena surrounded by display screens. **D**. Top view of the arena indicating the circular swimming area and preference sectors oriented toward each display screen. **E**. Experimental design. Focal fish were presented with two primary shoals and, in trichotomous trials, an additional decoy shoal. Trials were conducted in two order conditions: **(i)** dichotomous-first and **(ii)** trichotomous-first. Each trial began with a 2 min blank screen, followed by two 3 min stimulus presentation phases separated by a 2 min blank screen. In the dichotomous-first condition the two-option choice was presented first and the three-option choice second, whereas this order was reversed in the trichotomous-first condition. The positions of the primary shoals were counterbalanced within trials to avoid side bias, and the decoy position was counterbalanced across trials.

The main tank was elevated on a raised platform so that each face of the tank exactly matched the screen dimensions. The cylindrical design of the focal tank enabled the presentation of stimulus shoals from all four surrounding display screens while reducing corner effects common in rectangular or square choice tanks. Potential optical refraction from the curved surface of the focal tank was minimized by maintaining a water-filled space between the focal tank and the edges of the glass tank. The design was adapted from our previous studies using the cylindrical shoaling task for multi-alternative shoal-size preference assays (Singh et al., 2026).

Behavioural trials were recorded using a HP w300 webcam (1080p, 30 fps) mounted on a tripod directly above the setup. The entire setup was enclosed within a 3 m x 3 m black photography cloth to minimize external visual distractions, ensuring that the only light sources were the display screens.

### Virtual shoal movies

To generate virtual shoal movies, live fish were first recorded in a 3.5-L cuboidal tank (28 × 11 × 18 cm, L × B × H). Shoals of four individuals were sourced from a population tank containing mixed-sex adult fish, and a total of 10 such shoals were recorded 5 min each, using a GoPro Hero 8 camera (30 fps, 1080p, linear). The camera was positioned at a fixed distance from the tank such that the tank occupied the entire camera frame. Each shoal was allowed to acclimate for 5 minutes prior to recording. All sides of the recording tank, except the camera-facing side, were surrounded with black paper to avoid any visual distraction.

For each shoal, a 30 sec clip was selected in which individuals spent minimal time near the edges or the water surface. These clips were analyzed using multi-animal DeepLabCut pose-estimation software, with head, body, and tail points defined for each individual to form a skeletal representation for tracking movement. As the ‘body’ point was the most reliably tracked across individuals, ‘body’ point trajectories were subsequently used to generate virtual shoals using a custom R script.

Trajectories derived from the four-individual shoal video clips were used to construct virtual shoals movies of different shoal sizes (1, 2, 4, and 6 individuals). This ensured that differences in movement kinematics were not confounded by group size, other than the intended manipulation of shoal size itself.

Virtual shoal stimulus movies were generated in two steps. In the first step, a custom R script (R v 2022.07.2) was used to process cleaned coordinate datasets obtained from the multi-animal DeepLabCut, a Python-based pose estimation package (Lauer et al., 2022), analysis and to animate fish as black dots against a white background. Shoal size was manipulated by varying the number of individual trajectories included to form a virtual shoal, with each dot representing an individual fish. Dot size was adjusted such that the dot measured 1 cm in diameter on the display screens.

In the second step, all animated video files were imported into the video editing software Wondershare Filmora X (v 10.0.10.10.20) to control the remaining experimental variables: speed and spatial spread. Shoal speed was manipulated by altering the playback speed of the animation videos. Spatial spread was controlled by adjusting the proportion of the screen area occupied by the shoal, which was maintained at a centered 65% of the screen area across all virtual shoal movies. A five minute virtual shoal movie was generated by randomly assorting the animated 30 sec clips for each condition defined for the given experiment.

### Experimental design

#### Experiment 1

In the first experiment, we tested male and female subjects using dichotomous contrasts of virtual shoals that differed in both speed and shoal size, to establish a no-baseline preference condition before testing for decoy effects (Figure 2 A). In an exploratory manner, we tested two contrasts: 4 fish at 0.5x speed vs. 2 fish at 1.25x speed, and 4 fish at 0.75x speed vs. 2 fish at 1.25x speed.

**Figure 2.**
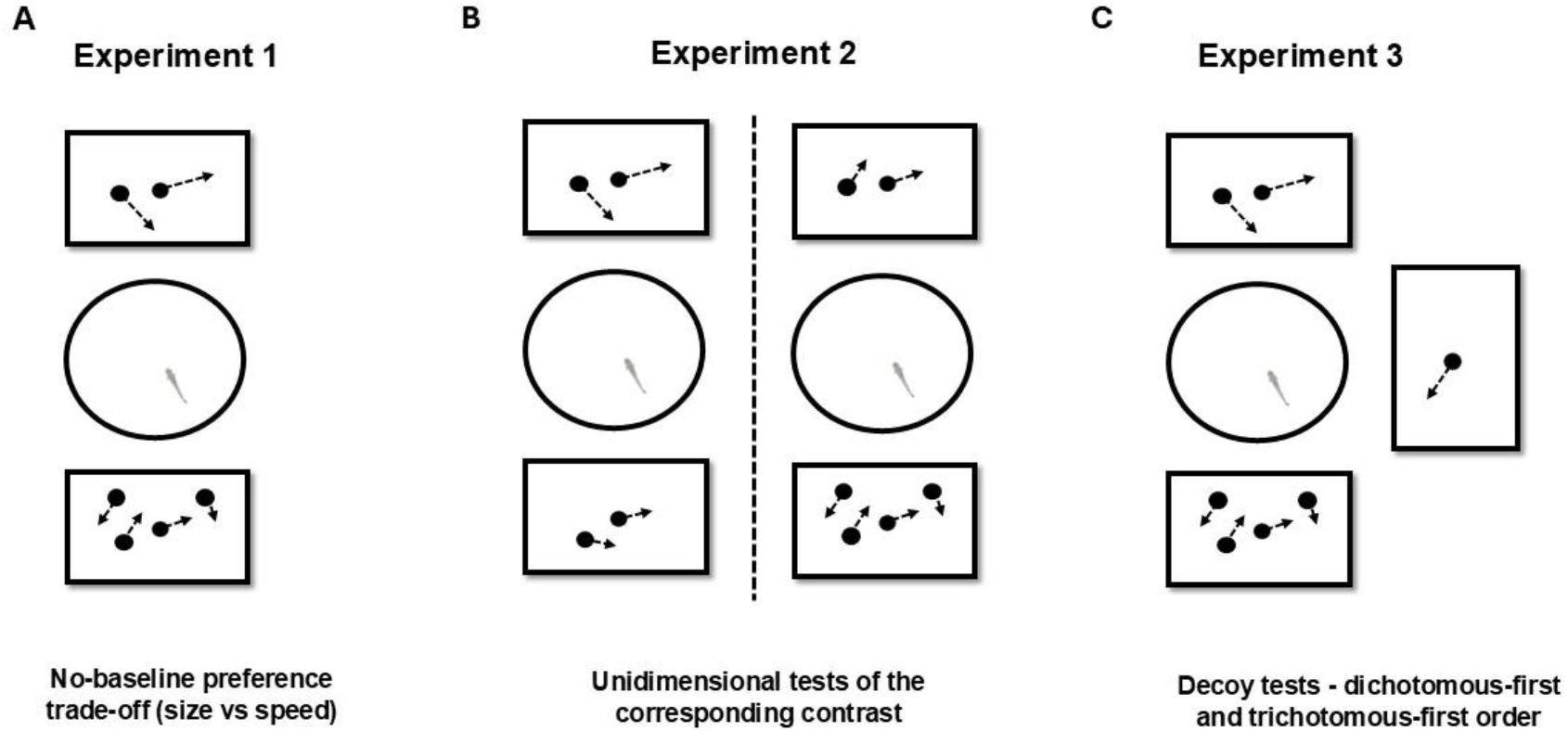
Experimental contrasts across Experiments 1–3. **A.** Experiment 1: Shoal preference tests contrasting two dimensions of size and speed, in a trade-off design: a larger, slower shoal (4 fish) versus a smaller, faster shoal (2 fish), establishing a no-baseline preference contrast. **B**. Experiment 2: Using the contrast that showed no baseline preference in Experiment 1, shoal preferences were tested by varying one dimension (size or speed) while holding the other constant, to determine whether equal weighting of both cues produced indifference in the two-dimensional trade-off. **C**. Experiment 3: For the same a no-baseline preference contrast, shoal preferences were tested by introducing a decoy shoal asymmetrically positioned on one of the attribute dimensions (size or speed), to examine context-dependent shifts in shoal preference.

In each dichotomous trial, subjects were introduced into the tank and first presented with 2 minutes of blank white screens to acclimatize them to the setup. This was followed by a 3-minute presentation of the two shoals on two of the four screens placed opposite to each other. The positions of the shoals were counterbalanced between trials. A total of 36 males and 36 females were tested in Experiment 1, with 18 males and 18 females assigned to each contrast.

#### Experiment 2

In experiment 2, for the contrast that had no baseline preference for either of the shoal in Experiment 1, we tested in unidimensional conditions, varying either shoal size or speed while keeping the other dimension constant, for both sexes (Figure 2 B). For the contrast 4 fish at 0.75x speed vs. 2 fish at 1.25x speed contrast, we performed (i) shoal size tests at constant speeds: 4 vs. 2 at 0.75x speed and 4 vs. 2 at 1.25x speed, and (ii) speed tests at constant shoal sizes: 0.75x vs. 1.25x at shoal size 4 and 0.75x vs. 1.25x at shoal size 2. A total of 73 males and 73 females were tested in Experiment 2, with 18–19 individuals of each sex assigned to each contrast.

#### Experiment 3

Two criteria were used to select the primary shoal contrast condition for Experiment 3. Based on the results of Experiment 1 and 2, the chosen contrast had to (i) show no baseline preference when shoals varied simultaneously in both speed and size, and (ii) show a preference for the larger shoal and the faster-moving shoal in the corresponding unidimensional tasks. This ensured that the indifference between primary shoals in the two-dimensional task reflected integration of information across both dimensions rather than a lack of sensitivity to either dimension.

Based on these criteria, the contrast 4 fish at 0.75x speed vs. 2 fish at 1.25x speed was selected. We then tested for the asymmetric dominance (decoy) effect by strategically placing decoy options that were extreme on either of the two dimensions (Figure 2 C). Specifically, we presented two shoal size decoys (6 fish at 0.75x and 1 fish at 0.75x) and two shoal speed decoys (2 fish at 0.5x and 4 fish at 1.5x).

For each decoy condition, shoals were tested under two order conditions: dichotomous-first and trichotomous-first. The primary shoal options were always presented on screens facing opposite each other. During trichotomous trials, the decoy option was displayed on one of the two remaining screens, while the fourth screen showed a blank white background. The positions of shoals in the dichotomous contrast were counterbalanced between dichotomous and trichotomous presentation within each trial, and the screen used for decoy presentation was counterbalanced between trials. A total of 201 males were tested in Experiment 3, with 23–29 individuals assigned to each contrast.

In a typical dichotomous-first trial, the subject was introduced into the tank and first presented with 2 minutes of blank screens, followed by a 3-minute presentation of the dichotomous contrast. This was followed by another 2-minute blank period and then a 3-minute presentation of the trichotomous condition including the decoy. The shoaling behavior was recorded only during the 3-minute shoal presentation phase.

### Behavioural coding

Each shoal preference recording was analysed using the behavioral coding software BORIS (v. 8.27.9) to quantify the time spent in each of the two sectors facing the stimulus screens in the dichotomous trials of Experiments 1 and 2, and in each of the three sectors facing the stimulus screens in the trichotomous trials of Experiment 2. Two investigators independently coded each video, and the recordings were verified for consistency between coders for each preference sector.

### Analysis

The circular arena was divided into two opposite angular sectors, facing the screens with primary shoals. Each sector had a radius of 25 cm and spanned 90° and served as a preference zone. for assessing shoaling behaviour (Figure 1 C). Time-spent data from the BORIS analysis were used to calculate the preference index for the target shoal based on the time spent in each sector, as follows:

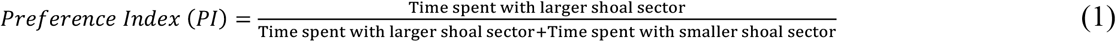

One-sample t-tests were used to determine whether the relative preference for the target shoal deviated from chance (preference index of 0.5) for each of the contrasts in all experiments. In Experiment 3 only the time spent in the sectors facing the primary shoal contrast was included to calculate the preference index for the target shoal. Since each test addressed independent and parallel hypotheses without influencing one another, the alpha level for all tests was maintained at 0.05.

## Results

### Experiment 1

To establish a no-baseline preference between shoals differing simultaneously in size and speed, we tested the dichotomous contrast between 4 fish at 0.5x speed and 2 fish at 1.25x speed. Both sexes deviated significantly from chance, preferring the 2 fish at 1.25x shoal (Figure 3, Table 1: Males, Mean = 0.25, SEM = 0.04, t(17) = −4.79, p < 0.01; Females, Mean = 0.42, SEM = 0.03, t(17) = −2.54, p < 0.05).

**Table 1.** Results from one-sample t-test for preference index comparisons against chance for Experiment 1, 2 and 3 for each shoal contrast conditions.

|  | Mean | +/- SEM | t | p |
| --- | --- | --- | --- | --- |
| **Experiment 1** |  |  |  |  |
| **4 fish 0.5x vs 2 fish 1.25x** |  |  |  |  |
| Males (N=18) | 0.25 | 0.04 | -4.79 | **<0.01\*\*** |
| Females (N=18) | 0.42 | 0.03 | -2.54 | **0.021\*** |
| **4 fish 0.75x vs 2 fish 1.25x** |  |  |  |  |
| Males (N=18) | 0.52 | 0.04 | 0.46 | 0.64 |
| Females (N=18) | 0.52 | 0.05 | 0.43 | 0.66 |
| **Experiment 2** |  |  |  |  |
| **4 fish 0.75x vs 2 fish 0.75x** |  |  |  |  |
| Males (N=18) | 0.56 | 0.02 | 2.19 | **0.04\*** |
| Females (N=18) | 0.65 | 0.04 | 3.16 | **<0.01\*\*** |
| **4 fish 1.25x vs 2 fish 1.25x** |  |  |  |  |
| Males (N=18) | 0.64 | 0.03 | 5.01 | **<0.01\*\*** |
| Females (N=18) | 0.65 | 0.04 | 3.07 | **<0.01\*** |
| **4 fish 1.25x vs 4 fish 0.75x** |  |  |  |  |
| Males (N=19) | 0.66 | 0.03 | 4.74 | **<0.01\*** |
| Females (N=19) | 0.57 | 0.06 | 1.14 | 0.26 |
| **2 fish 1.25x vs 2 fish 0.75x** |  |  |  |  |
| Males (N=18) | 0.58 | 0.03 | 2.70 | **0.01\*** |
| Females (N=18) | 0.58 | 0.04 | 1.66 | 0.11 |
| **Experiment 3** |  |  |  |  |
| **4 fish 0.75x vs 2 fish 1.25x \| Males \| Dichotomous-first** |  |  |  |  |
| **Decoy – 1 fish 0.75x (N=29)** |  |  |  |  |
| Dichotomous | 0.5 | 0.04 | 0.03 | 0.97 |
| Trichotomous | 0.46 | 0.03 | -1.24 | 0.22 |
| **Decoy – 2 fish 0.5x (N=29)** |  |  |  |  |
| Dichotomous | 0.58 | 0.03 | 2.09 | 0.05 |
| Trichotomous | 0.48 | 0.03 | -0.45 | 0.64 |
| **Decoy – 4 fish 1.5x (N=24)** |  |  |  |  |
| Dichotomous | 0.52 | 0.03 | 0.51 | 0.61 |
| Trichotomous | 0.51 | 0.02 | 0.58 | 0.56 |
| **Decoy – 6 fish 0.75x (N=24)** |  |  |  |  |
| Dichotomous | 0.51 | 0.04 | 0.22 | 0.82 |
| Trichotomous | 0.51 | 0.03 | 0.19 | 0.84 |
| **4 fish 0.75x vs 2 fish 1.25x \| Males \| Trichotomous-first** |  |  |  |  |
| **Decoy – 1 fish 0.75x (N=24)** |  |  |  |  |
| Trichotomous | 0.54 | 0.04 | 0.77 | 0.44 |
| Dichotomous | 0.58 | 0.04 | 1.69 | 0.10 |
| **Decoy – 2 fish 0.5x (N=24)** |  |  |  |  |
| Trichotomous | 0.49 | 0.03 | -0.19 | 0.84 |
| Dichotomous | 0.52 | 0.02 | 0.80 | 0.43 |
| **Decoy – 4 fish 1.5x (N=23)** |  |  |  |  |
| Trichotomous | 0.49 | 0.03 | -0.38 | 0.70 |
| Dichotomous | 0.47 | 0.04 | -0.60 | 0.54 |
| **Decoy – 6 fish 0.75x (N=24)** |  |  |  |  |
| Trichotomous | 0.52 | 0.03 | 0.59 | 0.55 |
| Dichotomous | 0.5 | 0.03 | -0.36 | 0.71 |

\*p<0.05 \*\*p<0.01

**Figure 3.**
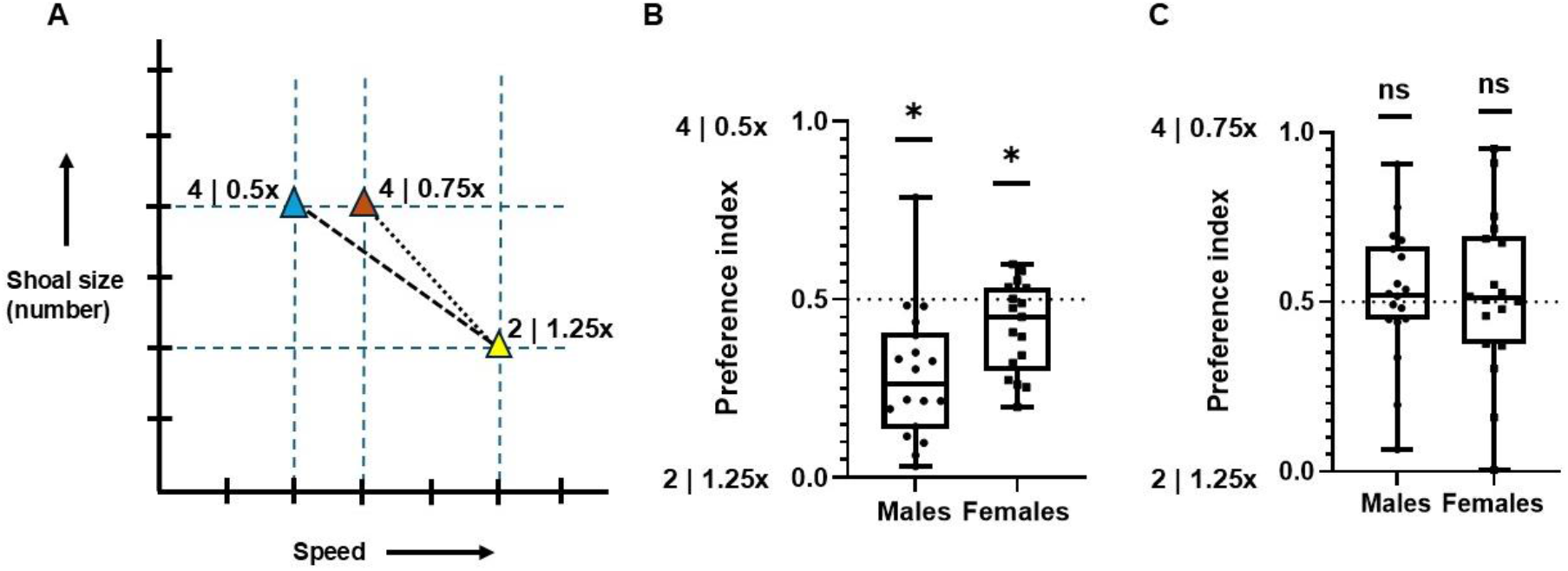
Experiment 1: Baseline dichotomous shoal preference test. **A.** Schematic representation of the two-dimensional choice space defined by shoal size (number of individuals) and speed. The contrasts compared 4 fish at 0.5x speed versus 2 fish at 1.25x speed, and 4 fish at 0.75x versus 2 fish at 1.25x speed. **B**. Preference indices for the contrast 4 fish at 0.5x vs 2 fish at 1.25x, shown separately for males and females. Both sexes showed a significant preference for 2 fish at 1.25x speed relative to chance (0.5; p < 0.05). **C**. Preference indices for the contrast 4 fish at 0.75x vs 2 fish at 1.25x. Neither males nor females differed significantly from chance (0.5; p > 0.05), indicating no baseline preference.

When presented with the dichotomous contrast between 4 fish at 0.75x speed and 2 fish at 1.25x speed, neither sex deviated significantly from chance (Figure 3, Table 1: Males, Mean = 0.52, SEM = 0.04, t(17) = 0.46, p >0.05; Females, Mean = 0.52, SEM = 0.05, t(17) = 0.43, p > 0.05). Thus, this contrast showed no baseline preference and was selected for subsequent experiments.

### Experiment 2

We next examined whether subjects discriminated shoal size independently of speed using unidimensional contrasts derived from the baseline-neutral condition. When shoal size varied (4 vs. 2) while speed was held constant at 0.75x, both sexes showed a significant preference for the larger shoal (Figure 4, Table 1: Males, Mean = 0.56, SEM = 0.02, t(17) = 2.19, p < 0.05; Females, Mean = 0.65, SEM = 0.04, t(18) = 3.16, p < 0.01). A similar pattern was observed when speed was held constant at 1.25x, with both sexes again preferring the larger shoal (Figure 4, Table 1: Males, Mean = 0.64, SEM = 0.03, t(17) = 5.01, p < 0.01; Females, Mean = 0.65, SEM = 0.04, t(18) = 3.07, p < 0.01).

**Figure 4.**
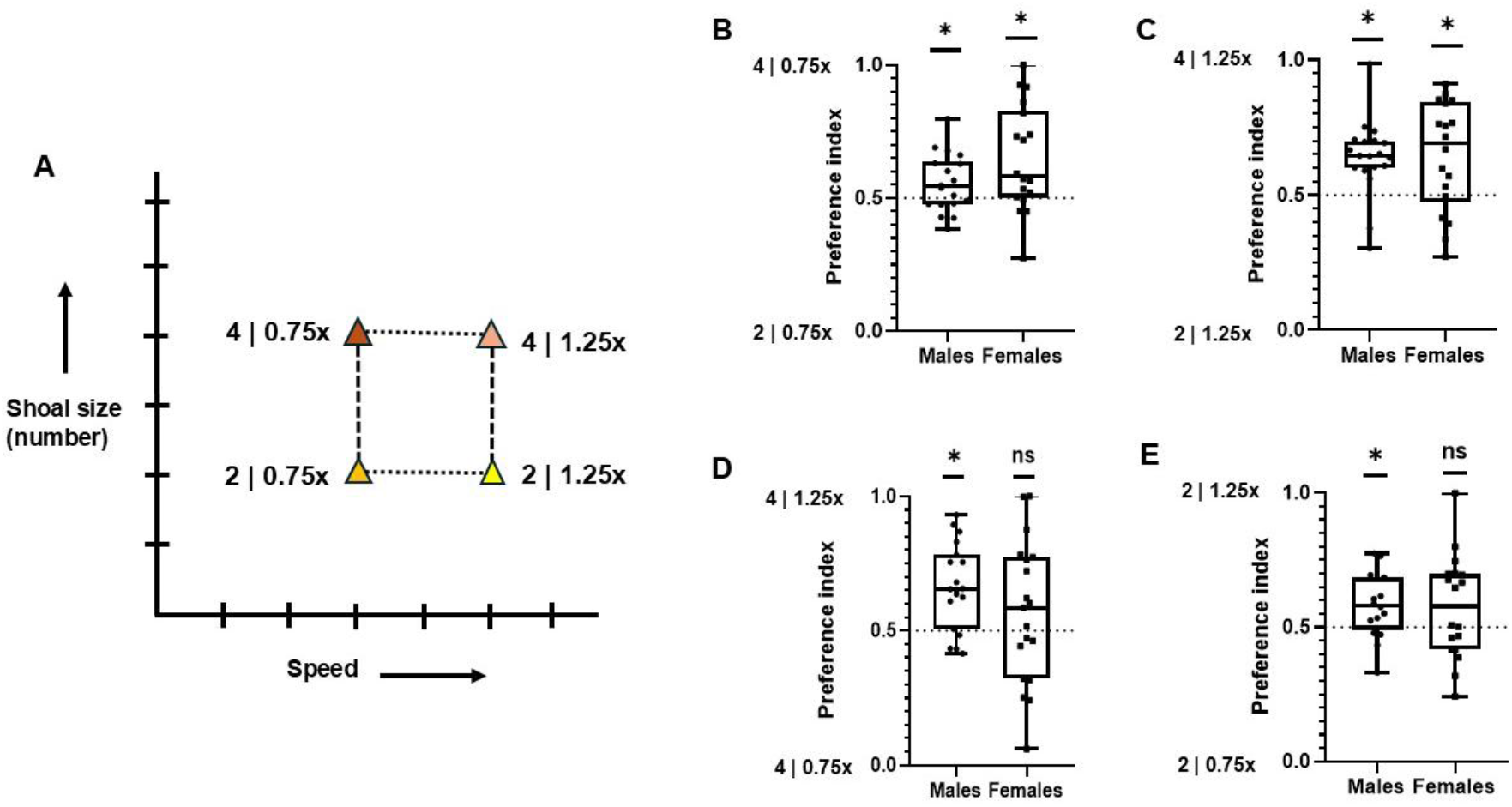
Experiment 2: Unidimensional shoal preference tests. **A.** Two-dimensional choice space illustrating contrasts in which one dimension (shoal size or speed) was varied while the other was held constant. Horizontal contrasts represent speed differences at constant shoal size (4 fish: 0.75x vs 1.25x; 2 fish: 0.75x vs 1.25x), whereas vertical contrasts represent size differences at constant speed (0.75x: 4 vs 2 fish; 1.25x: 4 vs 2 fish). **B–E**. Preference indices for each unidimensional contrast, shown separately for males and females. **B**. 4 fish at 0.75x vs 2 fish at 0.75x; both males and females preferred 4 fish at 0.75x significantly against chance (0.5; p < 0.05). **C**. 4 fish at 1.25x vs 2 fish at 1.25x; both males and females preferred 4 fish at 1.25x significantly against chance (0.5; p < 0.05). **D**. 4 fish at 1.25x vs 4 fish at 0.75x; males preferred 4 fish at 1.25x significantly against chance (0.5; p < 0.05) while females were indifferent (0.5; p > 0.05). **E**. 2 fish at 1.25x vs 2 fish at 0.75x; males preferred 2 fish at 1.25x significantly against chance (0.5; p < 0.05) while females were indifferent (0.5; p > 0.05).

When speed varied (1.25x vs. 0.75x) while shoal size was held constant at four fish, males showed a significant preference for the faster shoal, whereas females did not differ from chance (Figure 4, Table 1: Males, Mean = 0.66, SEM = 0.03, t(18) = 4.74, p < 0.01; Females, Mean = 0.57, SEM = 0.06, t(18) = 1.14, p > 0.05). A similar pattern was observed at shoal size two, with males again preferring the faster shoal, while females showed no significant preference (Figure 4, Table 1: Males, Mean = 0.58, SEM = 0.03, t(17) = 2.70, p < 0.05; Females, Mean = 0.58, SEM = 0.04, t(18) = 1.66, p > 0.05).

These results indicate that both males and females consistently preferred larger shoals across speeds, whereas preference for faster-moving shoals was evident only in males at both shoal sizes.

### Experiment 3

Based on the absence of baseline preference in Experiment 1 and the clear unidimensional discrimination of both shoal size and speed in males in Experiment 2, the contrast between 4 fish at 0.75x speed and 2 fish at 1.25x speed was selected to test for asymmetric dominance effects in males.

We found no evidence of an asymmetric dominance effect under either the dichotomous-first or trichotomous-first order of presentation. Within each order condition, preference indices did not deviate significantly from chance in either the dichotomous or trichotomous treatments for any decoy type (all p > 0.05; see Figure 5, Table 1).

**Figure 5.**
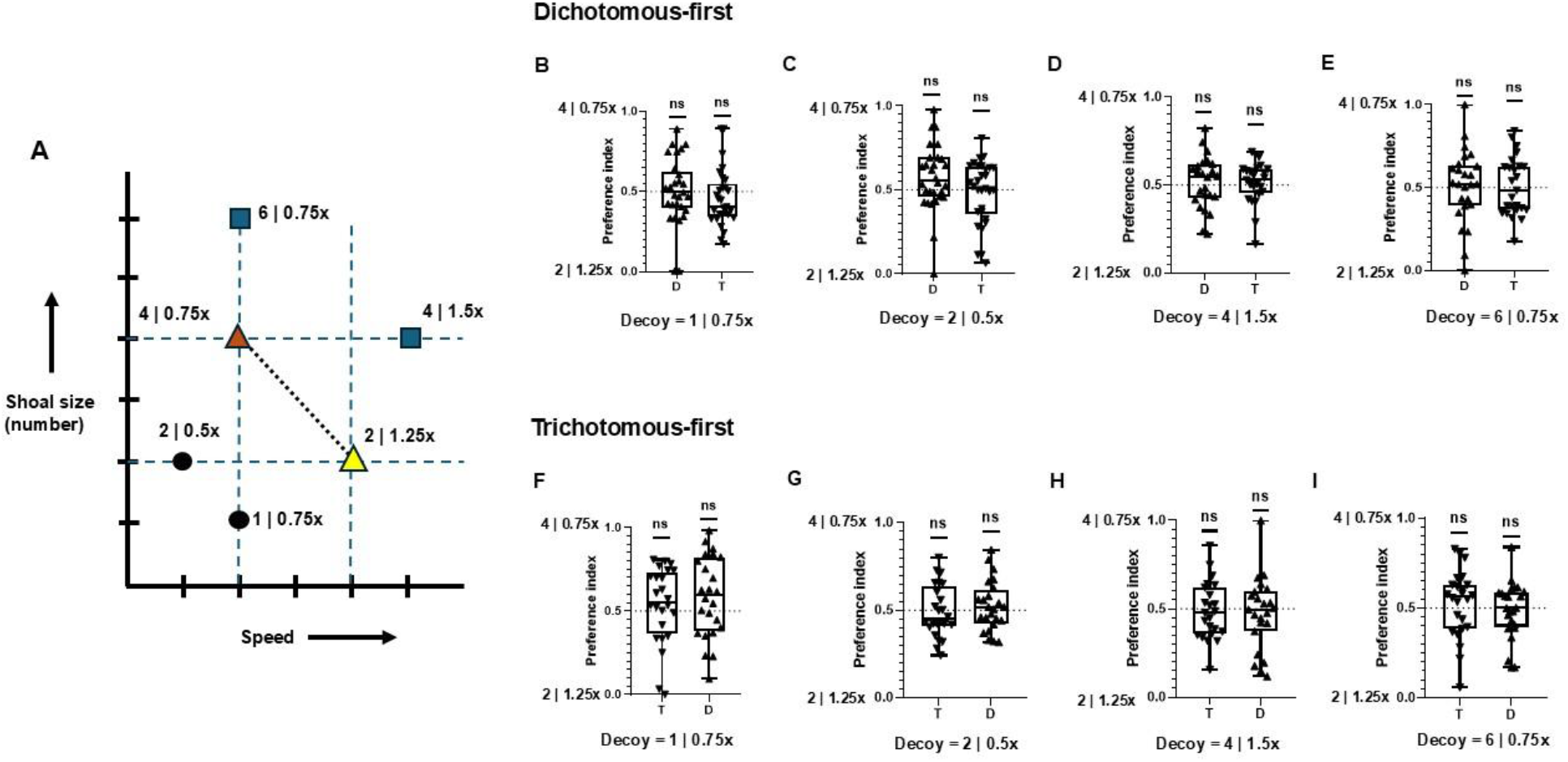
Experiment 3: Decoy effects under different order conditions. **A.** Two-dimensional choice space defined by shoal size (number of individuals) and speed, illustrating the primary trade-off contrast (4 fish at 0.75x vs 2 fish at 1.25x) and the positions of asymmetrically dominated decoys (1 fish at 0.75x, 2 fish at 0.5x, 4 fish at 1.5x, and 6 fish at 0.75x). **B–E**. Preference indices for the primary contrast under the dichotomous-first order condition. **F–I**. Preference indices for the primary contrast under the trichotomous-first order condition. Each panel corresponds to a different decoy option. No significant deviation from chance (0.5; p > 0.05) was detected in any decoy or order condition.

## Discussion

We tested whether zebrafish integrate information about shoal size and movement speed during choice, and whether such decisions are influenced by the contextual introduction of an asymmetrically placed decoy option. When shoals differed simultaneously in size and speed (4 fish at 0.75x vs. 2 fish at 1.25x), both sexes showed no baseline preference. In unidimensional tests, both males and females discriminated shoal size at both speeds, whereas discrimination of speed was observed only in males. Notably, the size preference in males was stronger when shoals moved faster. The introduction of asymmetric decoy options did not shift the relative preferences of male zebrafish between the primary shoals under either order of presentation.

### Multi-attribute integration and sex-specific perceptual weighting in zebrafish shoal choice

When presented with a larger but slower shoal (4 fish moving at 0.75x speed) versus a smaller but faster shoal (2 fish moving at 1.25x speed), both male and female zebrafish distributed their time equally between the two options. This suggests that the shoal attractiveness conferred by shoal size and that conferred by movement speed can be rendered approximately equivalent in subjective value. In an earlier contrast—4 fish moving at 0.5x speed versus 2 fish moving at 1.25x speed—both sexes showed a clear preference for the smaller, faster shoal. The fact that increasing the speed of the larger shoal from 0.5x to 0.75x was sufficient to eliminate this preference suggests that zebrafish are sensitive to graded differences in movement speed and can weigh these against numerical (shoal size) cues. This interpretation is consistent with evidence that fish shoaling decisions are not governed by a single dominant cue but instead arise from an integrated assessment of multiple ecologically relevant attributes (Swaney et al., 2025b; Ward et al., 2020). The role of movement speed as a weighted social cue in zebrafish is further supported by Lemasson et al. (2018), who showed that zebrafish consistently track virtual conspecific silhouettes moving faster than the background regardless of their number or directional coherence, whereas silhouettes moving at background speed elicit no consistent response. Movement speed therefore appears to be actively weighted when zebrafish evaluate social options.

Both males and females reached indifference at the same trade-off point. This contrasts with the pattern observed in the unidimensional experiments, where sex differences in speed sensitivity were apparent. Females showed reliable size preferences when speed was held constant, yet this preference was disrupted when speed and size were traded off, despite speed carrying no independent weight for females. Similar interference of movement speed with numerical discrimination of shoal size was reported by Agrillo et al. (2008), in which female mosquitofish failed to discriminate between shoals of 4 vs. 8 female conspecifics when the activity levels of the shoals were matched by increasing the temperature of the smaller shoal and decreasing that of the larger shoal. This suggests that variation along an otherwise irrelevant dimension may introduce sufficient perceptual interference to disrupt size-based discrimination, rather than reflecting a genuine trade-off between two weighted attributes. This pattern is consistent with the logic of Garner interference: varying a task-irrelevant dimension orthogonally to a task-relevant one can impair discrimination along the relevant dimension, even when the irrelevant dimension exerts no independent influence on choice (Algom & Fitousi, 2016).

The present study revealed a clear sexual dimorphism in speed preference: male zebrafish preferred the faster-moving shoal at both shoal sizes tested (2 vs. 2 and 4 vs. 4), whereas females showed no significant preference for speed at either size. Sex differences in the perceptual weighting of shoal attributes are not without precedent in zebrafish. Engeszer et al. (2008) demonstrated that males and females occupy fundamentally different perceptual spaces when assessing potential shoalmates, with males selectively responding to identifiable visual traits such as species identity and stripe patterning that females appear to disregard.

The present finding extends this principle from static visual features to dynamic ones, suggesting that sexually dimorphic perceptual filtering of shoal attributes may be a broader feature of zebrafish social cognition. The observed sex difference in speed sensitivity may reflect divergent predator-avoidance strategies, with males potentially interpreting faster movement as a signal of heightened vigilance or escape readiness in conspecifics, whereas females may rely more heavily on shoal size alone as a cue of safety.

When shoal size was the only varying attribute, both males and females preferred the larger shoal against chance regardless of background speed (0.75x or 1.25x). However, males exhibited only a weak preference at the lower playback speed (0.75x), which became stronger at the higher speed (1.25x). This may reflect speed-dependent modulation of attentional engagement: faster-moving stimuli are known to capture attention more strongly in zebrafish (Lemasson et al., 2018), and Puy et al. (2024) have proposed that fish selectively attend to faster-moving neighbours through a speed-based attentional mechanism. This pattern is also consistent with previous reports of sex differences in shoal size preference in zebrafish (Etinger et al., 2009; Ruhl & McRobert, 2005; Singh et al., 2026; Velkey et al., 2022), where males are often found to exhibit weaker or more variable preferences for larger shoals. Because male preferences may integrate additional cues such as movement speed and stripe patterning, whereas females appear to rely more strongly on shoal size alone, male choice may be more sensitive to variation in experimental conditions.

### No evidence for the asymmetric dominance effect in zebrafish shoal choice

Despite meeting the conditions considered necessary for a valid test of the asymmetric dominance effect among males, including indifference between primary options, demonstrated sensitivity to each dimension in isolation, and asymmetrically positioned decoys spanning both attribute dimensions, no decoy shifted preferences away from chance. This null result is not without precedent. Several studies have reported asymmetric dominance effects in bees, hummingbirds, and grey jays (Bateson et al., 2002, 2003; Shafir et al., 2002), but null or reversed effects have been documented in fish, primates, and rodents (Latty & Beekman, 2011; Parrish et al., 2015; Rivalan et al., 2017), and the effect shows considerable interspecific and inter-task variability overall. Armand et al. (2025) have recently reviewed decoy effects across pollinator studies, concluded that clear evidence for attraction effects in insects is largely lacking. A closer inspection of positive reports in insects reveals that many do not constitute true attraction effects. In honeybees, apparent shifts in preference resulted either from direct selection of the decoy rather than increased attraction to the target (Shafir et al., 2002), or from a reduction in competitor choices without any corresponding increase in target preference (Tan et al., 2015). In bumblebees, a reported preference shift was more parsimoniously explained by incentive contrast, in which exposure to a lower-quality option lowers the acceptance threshold, rather than by genuine contextual reweighting of attributes (Hemingway et al., 2024).

Frederick et al. (2014), replicating 38 human studies on asymmetrically dominated decoys, found reliable attraction effects only when options were compared in an abstract space (numerically) than experienced directly, a condition that cannot be met in animal studies where choices are necessarily experiential. Spektor et al. (2021), proposed that attraction, compromise, and similarity effects are highly sensitive to seemingly minor features of experimental design, including spatial arrangement of options, attribute concreteness, and deliberation time, and that context effects reliably disappear or reverse when attributes are represented perceptually. In the present study, shoal size and movement speed were experienced directly as perceptual quantities rather than described symbolically, a condition that Spektor et al. identify as particularly unfavourable for the emergence of attraction effects. Indeed, rather than a reliable attraction effect, some experiential choice paradigms have documented a repulsion effect, in which an asymmetrically inferior decoy actually decreases preference for the target it most resembles (Parrish et al., 2024). That neither attraction nor repulsion was observed in the present study suggests that zebrafish shoal preferences may be largely insensitive to third-option context under the conditions tested, consistent with a decision architecture that prioritises absolute attribute values over relational comparisons within the choice set.

### Biological motion stimuli provide valid multi-attribute social signals for zebrafish

A methodological contribution of this study is the development of biological motion stimuli in which shoal size and swimming speed can be independently and systematically manipulated, allowing shoal size preferences to be tested beyond the traditional use of biological motion as a simple affiliative stimulus. Sociability, defined as the tendency to associate with a shoal, and the ability to discriminate between shoals of different sizes are thought to reflect distinct processes, with the latter involving numerical or quantity discrimination and not necessarily correlating with general sociability. The fact that zebrafish discriminated both shoal size and movement speed from point-light displays confirms that these stimuli preserve socially relevant information present in real shoals. This approach offers several advantages over live-fish paradigms. It eliminates confounds arising from inter-individual variation in color, size, sex, and behavior, allows precise parametric manipulation of kinematic variables, and enables exact replication across studies. The paradigm therefore provides a useful platform for future investigations of the perceptual and cognitive mechanisms underlying multi-attribute social decision-making in zebrafish, including tests involving additional stimulus dimensions such as spatial spread, directional coherence, or inter-individual distance within the shoal.

## Conclusion

The present findings indicate that zebrafish integrate multiple dynamic attributes when evaluating conspecific groups, but that the perceptual weighting of these attributes differs between the sexes. While males incorporated both shoal size and movement speed in their decisions, females relied primarily on shoal size, with variation in movement speed interfering with shoal size discrimination rather than contributing an independently weighted cue. Despite testing under conditions designed to favour the asymmetric dominance effect, the introduction of asymmetrically dominated decoys did not alter preferences, suggesting that zebrafish shoal choice may operate largely independently of third-option context with respect to the two attributes of shoal speed and size. Instead, decisions appear to rely primarily on the absolute values of socially relevant attributes rather than on relational comparisons within the choice set. These results highlight sex-specific mechanisms of cue integration in zebrafish social decision-making and suggest that differences in how individuals weight dynamic social information may influence how local interactions scale up to the collective organisation of shoaling groups.

